# The neuronal homeobox transcription factor HMX3 is a crucial vulnerability factor in MECOM-negative KMT2A::MLLT3 acute myelomonocytic leukemia

**DOI:** 10.1101/2023.11.07.565950

**Authors:** Saioa Arza-Apalategi, Branco M. H. Heuts, Saskia M. Bergevoet, Roos Meering, Daan Gilissen, Pascal W. T. C. Jansen, Anja Krippner-Heidenreich, Peter J. M. Valk, Michiel Vermeulen, Olaf Heidenreich, Torsten Haferlach, Joop H. Jansen, Joost H. A. Martens, Bert A. van der Reijden

## Abstract

The KMT2A::MLLT3 fusion protein causes acute myeloid leukemia (AML) by activating the oncogenic transcription factor MECOM. However, *MECOM* expression occurs in only half of the KMT2A::MLLT3 cases. By integrating gene expression and enhancer activity data from patient cells, we identified neuronal homeobox transcription factor *HMX3* as cell fate determining factor in *MECOM*-negative KMT2A::MLLT3 AML. *HMX3* expression associated with younger age and KMT2A-rearranged leukemia in large AML cohorts (p<0.002). *HMX3* was not expressed in other major genetic risk groups and healthy blood cells. Transcriptomic analyses revealed that *HMX3* drives cancer-associated *E2F*, *MYC* and cell cycle gene programs. Ectopic *HMX3* expression completely inhibited monocytic but not granulocytic colony formation of healthy CD34^+^ adult cells. Silencing of *HMX3* in KMT2A::MLLT3 AML cell lines and patient cells resulted in cell cycle arrest, monocytic differentiation, and apoptosis. Thus, HMX3 is a leukemia-specific vulnerability that enhances proliferation and blocks differentiation of MECOM-negative KMT2A::MLLT3 leukemia.

## Introduction

Acute myeloid leukemia (AML) exhibits clinical, cytogenetic, and molecular heterogeneity ^1,2^. The recurrent chromosomal translocation t(9;11)(p22;q23) occurs in ∼5% of adult and ∼9% of pediatric AML cases, resulting in the fusion of *KMT2A* (a.k.a *MLL1*) to its most frequent partner *MLLT3* (a.k.a. *AF9*) ^3–6^. The resulting KMT2A::MLLT3 fusion protein contributes to leukemia development by direct chromatin binding to activate oncogene expression ^7–11^. According to European LeukemiaNet (ELN) criteria, adult KMT2A::MLLT3 AML is classified as intermediate risk AML and needs to be treated accordingly ^12^. Others and we previously showed that KMT2A::MLLT3-driven leukemia is separated into two groups based on the absence or presence of expression of the transcription factor (TF) MECOM (a.k.a. EVI1). KMT2A::MLLT3 patients with MECOM overexpression (MECOM^+^) have a particularly poor outcome, while its absence (MECOM^−^) associated with favourable outcomes and a very distinct gene expression profile (GEP) ^13–17^. MECOM itself is an important oncogenic factor in the MECOM^+^ KMT2A::MLLT3 subgroup ^17–20^. Even for the KMT2A::MLLT3 group with favourable outcome, the treatment comes with significant short and long term toxicity/morbidity, and thus needs to be improved ^21,22^. A better understanding of the targets of KMT2A::MLLT3 that determine cell fate and drive AML pathobiology may identify new targets for less toxic therapeutic intervention.

Normally, hematopoietic cell fate is orchestrated by complex gene regulatory networks (GRN) centered around TFs. TF dysregulation due to mutations, or gene fusions (such as KMT2A::MLLT3) can lead to alterations in GRNs that may result in hematopoietic cell transformation ^23,24^. In addition, changes in the activity of non-mutated TFs play an important role in leukemia as well ^25^. TFs that dictate leukemia cell fate through altered activity can be identified because they predominantly bind gene enhancers. By predicting genome-wide TF binding profiles in different cell types using enhancer activity and TF binding motifs, and by integrating these inferred binding profiles with genome-wide gene expression data, cell type-specific GRN and key TFs controlling cell fates can be identified ^26,27^. Of all currently available bioinformatic algorithms, the Analysis Algorithm for Networks Specified by Enhancers (ANANSE) performs best in identifying cell fate determining TFs ^26^. While such bioinformatic approaches are very powerful, they need genome-wide gene expression and enhancer activity data as input. For the latter, a proxy for open chromatin or histone H3K27-acetylation data are often used. We recently adapted the ANANSE approach by using solely 5’ Cap Analysis of Gene Expression sequencing (CAGE-seq) data as input (ANANSE-CAGE) ^27^. The advantage of CAGE-seq is that both expression of protein coding genes based on unidirectional expression as well as enhancer activity based on bidirectional expression of enhancer RNAs (eRNAs) can be determined in one go. In the present study, we generated and interrogated CAGE-seq data from primary MECOM^+^ and MECOM^−^ KMT2A::MLLT3 cases using ANANSE-CAGE and predicted neuronal H6 family homeobox 3 (HMX3) as cell fate determining TF in MECOM^−^ KMT2A::MLLT3 AML. Subsequently, we determined that HMX3 activates oncogenic GRN in MECOM^−^ KMT2A::MLLT3 acute myelomonocytic leukemia cells and that these cells rely on HMX3 for their block in differentiation and proliferation.

## Results

### Identification of HMX3 as cell fate determining TF in MECOM-negative KMT2A::MLLT3 AML

To identify TFs that determine cell fate and pathobiology, we first performed CAGE-seq to identify gene expression and enhancer activity differences between MECOM^+^ and MECOM^−^ KMT2A::MLLT3 AML using primary patient samples (n = 3, each). A total of 26,118 unidirectional transcription start sites (TSSs) and 13,751 bidirectional transcription regions (eRNAs) were identified (Supplementary Table S1). Principal component analysis (PCA) using every unidirectional TSS cluster per gene—which represents a proxy for gene expression—clearly separated MECOM^−^ and MECOM^+^ samples on PC1, in line with previous reported gene expression differences between these two groups (Figure 1A) ^15,17^. Importantly, eRNA activity, which represents a proxy for active enhancers, also separated the two KMT2A::MLLT3 subgroups (PC2) (Figure 1B), suggesting that we can exploit this difference using our enhancer-controlled network-based TF prediction method (ANANSE-CAGE) ^27^. In doing so, we indeed identified MECOM as the most influential cell fate determining factor in MECOM^+^ KMT2A::MLLT3 AML. In addition, other major oncogenic hematopoietic transcription factors were predicted as important in MECOM^+^ KMT2A::MLLT3 AML, such as MYB, ERG, GATA2 and TCF4 (Figure 1C) ^28–31^. In MECOM^−^ KMT2A::MLLT3 AML, different TFs were identified on which leukemia cells may rely, such as E2Fs, GFI1, IRF8, MYC, and AHR (Figure 1D) ^32–35^. Notably, the neuronal TF HMX3, that has no described function in blood cell development, emerged as the most influential within this subgroup. ^36^.

**Figure 1.**
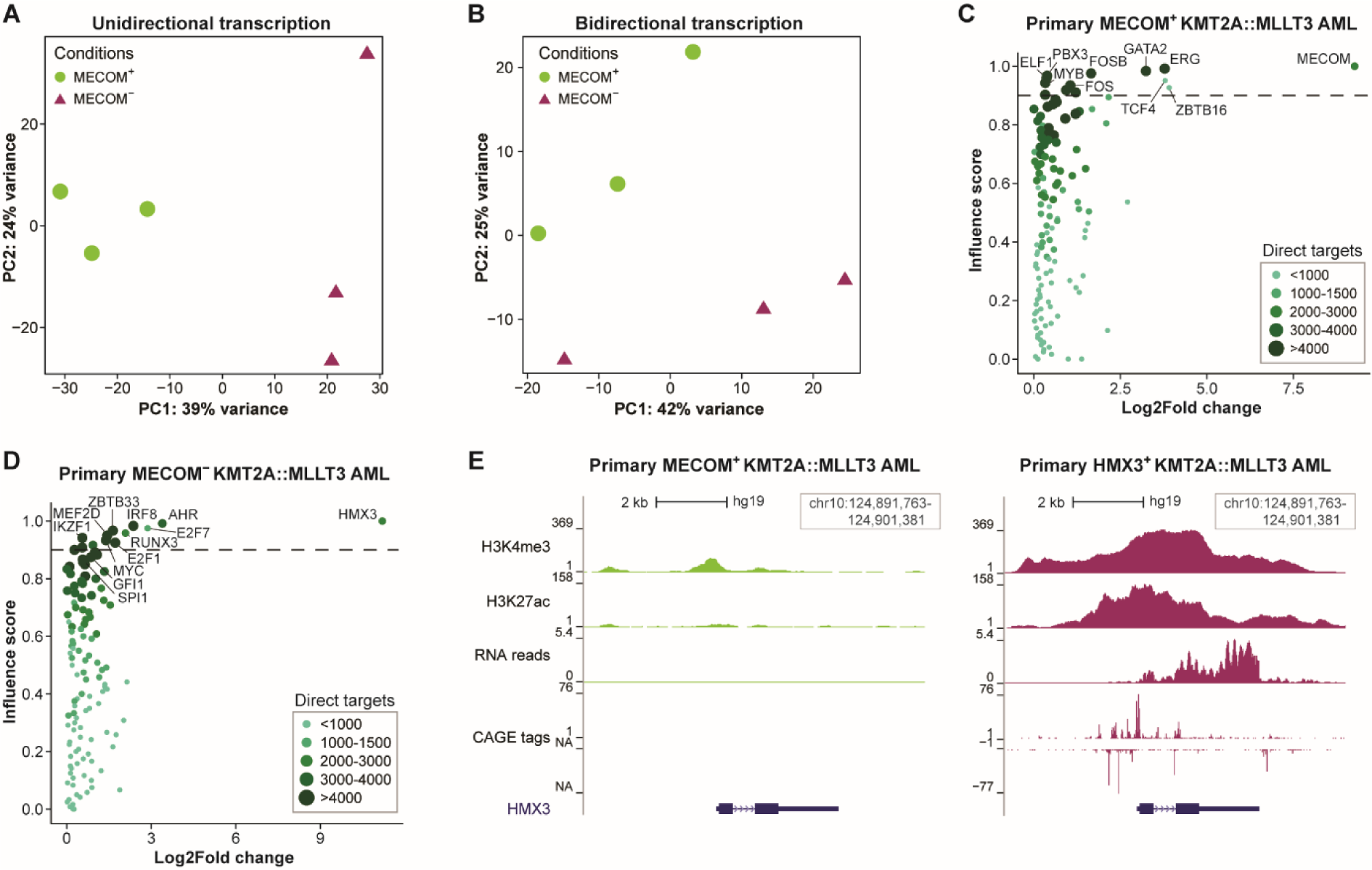
ANANSE identifies neuronal homeobox transcription factor HMX3 to determine MECOM^−^ KMT2A::MLLT3 AML cell fate. (**A**) PCA plot using 3 MECOM^+^ and 3 MECOM^−^ KMT2A::MLLT3 AML CAGE-seq samples (P1-6), part of which have been described in other studies ^8,37,51^. The sum of all unidirectional TSS clusters per gene is used as a proxy for gene expression. Colors and shapes represent the conditions. (**B**) Similar to (A), except using bidirectional transcription activity (enhancer activity). (**C**) Scatterplot depicting log2 fold change and influence scores. The influence score is an inferred score that describes how effectively differences between two cell states can be attributed to a TF. MECOM^+^ KMT2A::MLLT3 AML samples were compared to MECOM^−^ KMT2A::MLLT3 AML samples, thus revealing key TFs in MECOM^−^ KMT2A::MLLT3 AML. The color and size of the individual dots represent an approximation for the number of target genes that are calculated from the number of edges in differential GRNs. The dotted line represents a visual cut-off for highlighting the top ranked TFs. (**D**) Similar to (C), except MECOM^−^ KMT2A::MLLT3 AML samples were compared to MECOM^+^ KMT2A::MLLT3 AML samples, thus revealing key TFs in MECOM^+^ KMT2A::MLLT3 AML. (**E**) UCSC genome browser representation of the *HMX3* locus showing CAGE-, RNA-, and ChIP-seq (H3K27ac and H3K4me3) data from MECOM^+^ and MECOM^−^ KMT2A::MLLT3 AML patients, P3 and P6 respectively ^37^. The CAGE tags tracks are the result of merged CAGE tags from patients P1-P3 (MECOM^+^) and P4-6 (HMX3^+^). Colors match the conditions in (A).

To investigate epigenetic regulation of the *HMX3* locus, we analyzed H3K4me3 and H3K27ac ChIP-sequencing data from 2 KMT2A::MLLT3 AML samples (one of each subgroup) (Figure 1E) ^37^. This revealed that the *HMX3* locus is epigenetically very active in the MECOM^−^ HMX3-positive (HMX3^+^) case, while being epigenetically inactive in the MECOM^+^ case. Indeed, *MECOM* and *HMX3* are expressed in a mutually exclusive manner (Supplementary Figure S1A). In addition, by analyzing KMT2A::MLLT3 DNA binding from publicly available KMT2A::MLLT3-transduced CD34^+^ cord blood data ^11^, we observed *HMX3* as one of the 201 genomic regions that was bound by KMT2A::MLLT3 (Supplementary Figure S1B). *HMX3* was among the 62 early downregulated genes after KMT2A::MLLT3 degradation as determined by SLAM- and PRO-seq ^11^. Finally, we performed label free quantitative mass spectrometry on primary bone marrow from a MECOM^+^ and HMX3^+^ KMT2A::MLLT3 AML case (Supplementary Table S2). While peptides represented by MECOM but not HMX3 were readily detected in the MECOM^+^ case, the opposite was observed in the HMX3^+^ case. Together, these data corroborate our finding that HMX3 is key to MECOM^−^ KMT2A::MLLT3 AML.

### Distinct recurrent genetic abnormalities associate with HMX3 expression

Earlier, others and we showed that MECOM^−^ KMT2A::MLLT3 AML have a very distinct GEPs ^15,16^. To assess whether HMX3^+^ AML cases exhibit a similar GEP in general, we performed Uniform Manifold Approximation and Projection (UMAP) using the 2000 most variable protein coding genes from two AML cohorts, *i.e.* TARGET-AML (n = 178 pediatric cases) and Beat AML (n = 410 adult cases and 19 healthy bone marrow samples) (Figure 2A-B) ^38^. In both cohorts we observed a distinct cluster of HMX3^+^ cases, although we also identified a few HMX3^+^ cases that appeared to exhibit alternative GEPs. To unravel the genetics-based identity of HMX3^+^ cases, we examined the presence of recurrent genetic abnormalities (Figure 2C-D). This again showed that KMT2A::MLLT3 cases either express *MECOM* or *HMX3* in a mutually exclusive manner (Supplementary Figure S1C-D). In addition, we observed a strong correlation of *HMX3* expression with other *KMT2A*-rearranged cases, including the expected overrepresentation of HMX3^+^ KMT2A::MLLT3 AML cases in both cohorts (Supplementary Figure S1E-F). Two out of three cases positive for the t(8;16) resulting in the KAT6A::CREBBP fusion protein were also present in the main HMX3^+^ cluster in the Beat AML cohort. In contrast, *HMX3* was not expressed in most major genetically defined AML subgroups, such as CBFB::MYH11, RUNX1::RUNX1T1, or FLT3-ITD, as well as none of the healthy bone marrow samples (Figure 2E-F). Independent analyses of our earlier generated Blueprint consortium RNA-seq data confirmed that *HMX3* is not expressed in any of healthy blood cell types implying that *HMX3* expression is leukemia specific (Supplementary Figure S1G) ^39^. Overall, these results indicate that *HMX3* expression defines a distinct AML subtype characterized by a specific gene expression signature, suggesting a dominant role for HMX3 in leukemia-specific transcription regulation.

**Figure 2.**
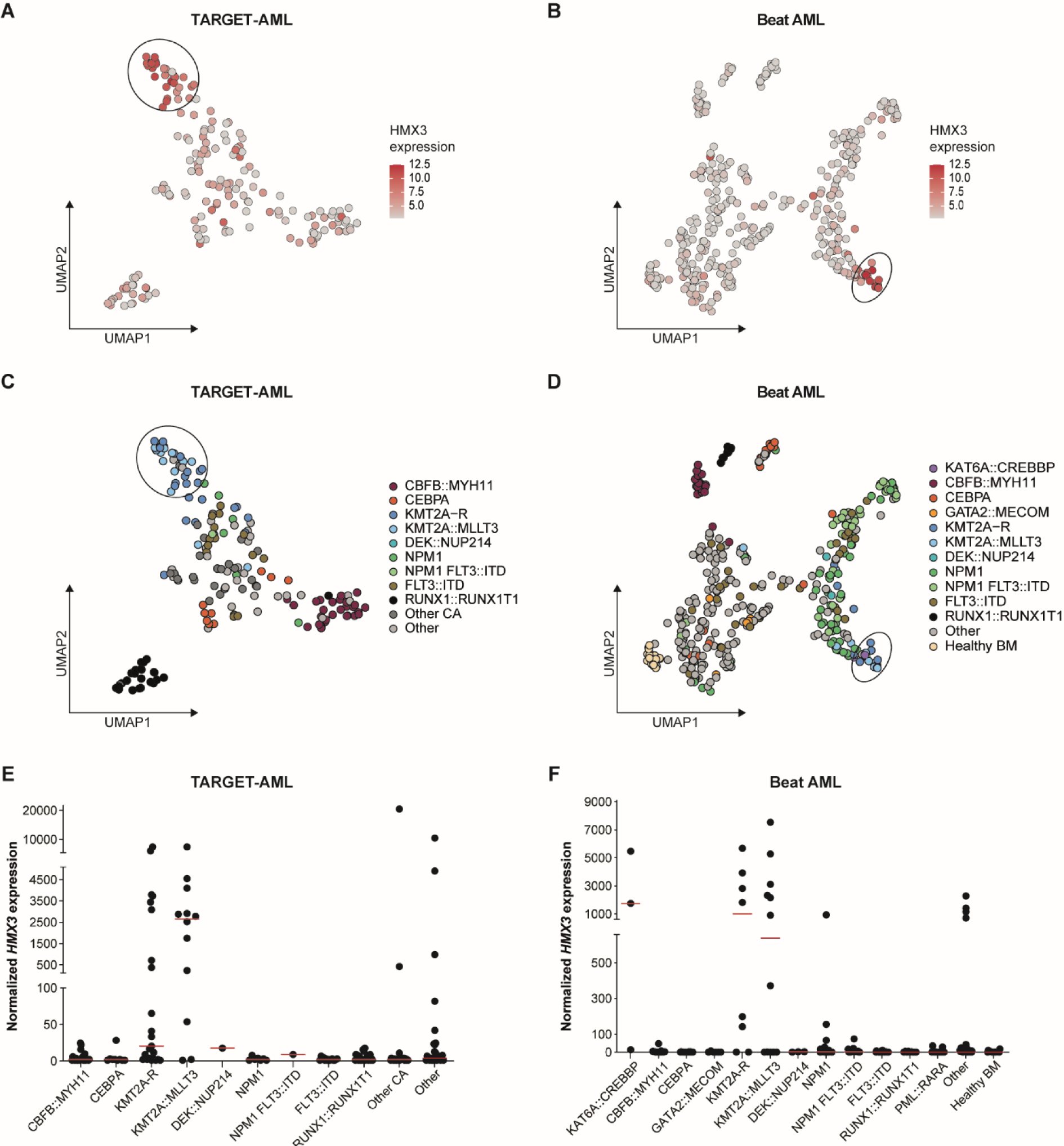
*HMX3* expression correlates with a distinct gene expression profiles of KMT2A-rearranged and t(8;16) AML. **(A)** Uniform Manifold Approximation and Projection (UMAP) was performed on the 2000 most variable protein-coding genes from patient samples of the TARGET-AML cohort. These samples were colored according to *HMX3* expression. The black circle highlights the *HMX3*+ patients that have a similar gene expression profile. **(B)** Similar to (A), except using the Beat AML cohort. **(C)** Similar to (A), except samples are colored by genetic abnormalities. ‘CA’ stands for cytogenetic abnormalities. **(D)** Similar to (C), except using the Beat AML cohort. **(E)** Distribution of DESeq2 normalized gene counts of *HMX3* in the TARGET-AML cohort per AML subtype. Red line shows the median value. **(F)** Similar to (E), except using the Beat AML study group.

### HMX3 regulates oncogenic gene programs in HMX3-positive KMT2A::MLLT3 AML

To uncover HMX3-dependent gene programs in KMT2A::MLLT3 AML, we first performed RNA sequencing using the HMX3^+^ KMT2A::MLLT3 cell line MOLM-13 and *HMX3* silenced (HMX3 KD) MOLM-13 (n=3 each). *HMX3* silencing was confirmed by qPCR (Supplementary Figure S1H). PCA showed a strong separation (91% variance) between the two conditions with 704 upregulated and 608 downregulated genes (padj < 0.05, respective fold change > 1.5 and < −1.5) following *HMX3* silencing (Figure 3A and Supplementary Table S3). To elucidate biological processes associated with *HMX3* expression, we performed Gene Set Enrichment Analysis (GSEA). This revealed significantly enriched gene sets related to active proliferation, such as E2F and MYC targets, and G2M checkpoint ^40–42^, in *HMX3* expressing cells (Figure 3B). Conversely, gene sets related to immune response, such as TNFA signaling, inflammatory response, IL6-JAK-STAT3 signaling, and interferon gamma response were significantly enriched following *HMX3* silencing (Figure 3B). To validate these findings, we performed differential expression analysis using primary MECOM^+^ and HMX3^+^ KMT2A::MLLT3 AML cases from the Beat AML cohort. Indeed, *HMX3* expressing KMT2A::MLLT3 cases showed enrichment of gene sets relevant for cell proliferation (E2F and MYC targets, and G2M checkpoint), whereas gene sets associated with immune response (TNFA signaling via NFKB, inflammatory response, IL6-JAK-STAT3 signaling, and interferon gamma response) were significantly enriched in the MECOM^+^ KMT2A::MLLT3 cases (Figure 3D-F). These results indicate that HMX3 regulates oncogenic gene programs.

**Figure 3.**
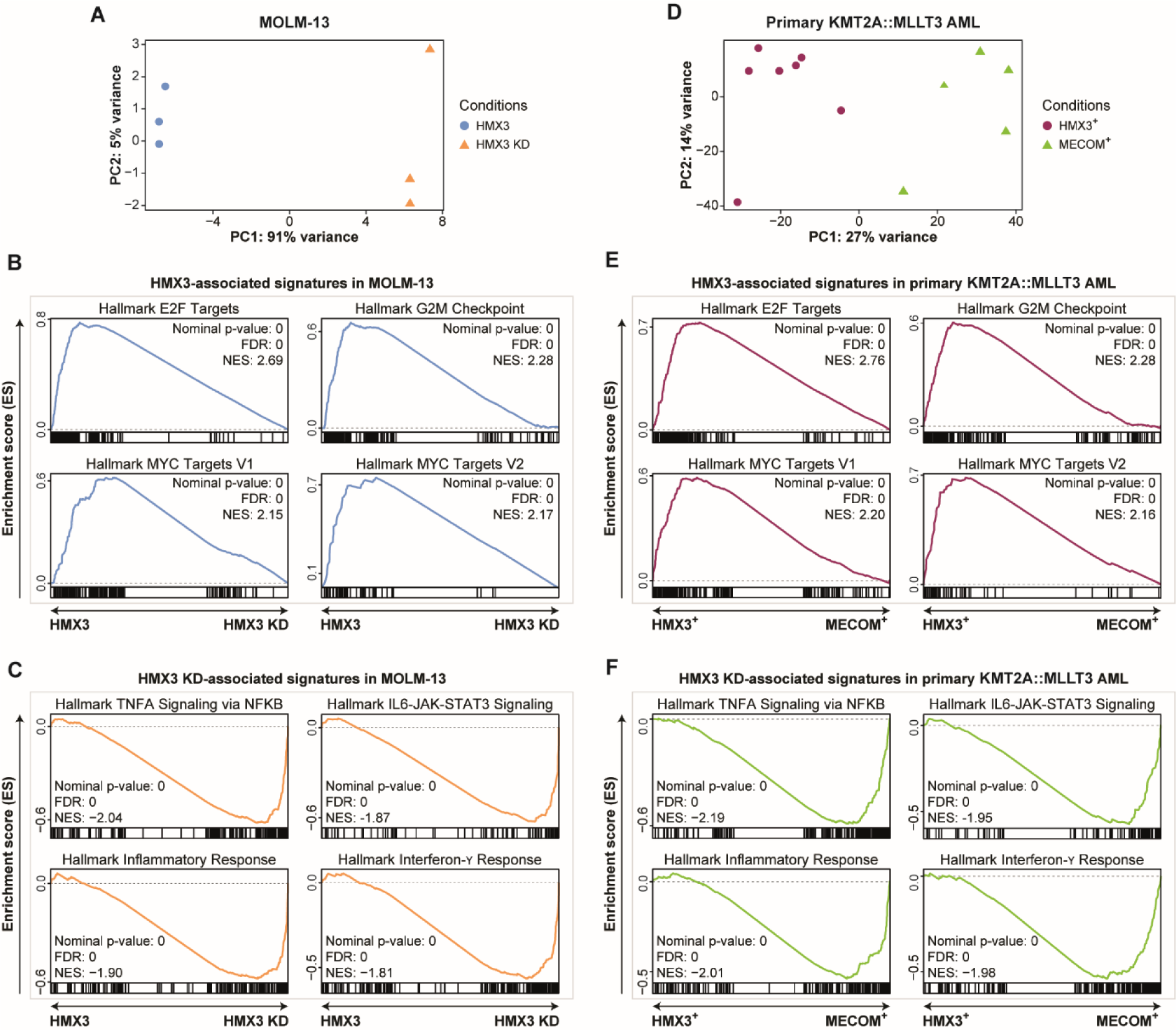
*HMX3* regulates E2F and MYC programs. **(A)** PCA plot using RNA-seq data from MOLM-13 cells transduced with shRNA against *HMX3* (HMX3 KD) or non-targeting control (*HMX3*). Colors and shapes represent the conditions**. (B)** Enriched gene sets in *HMX3*+ MOLM-13 cells that were transduced with non-targeting shRNA control (*HMX3*). Colors represent the conditions from (A). **(C)** Enriched gene sets in MOLM-13 cells that were transduced with shRNA against *HMX3* (HMX3 KD). Colors represent the conditions from (A). **(D)** Similar to (A), except using primary KMT2A::MLLT3 AML samples from the Beat AML cohort. **(E)** Enriched gene sets in HMX3+ KMT2A::MLLT3 AML (HMX3+). Colors represent the conditions from (D). **(F)** Enriched gene sets in MECOM+ KMT2A::MLLT3 AML (MECOM+). Colors represent the conditions from (D).

### HMX3 activates oncogenic gene programs in healthy CD34-positive cells and inhibits their clonogenic growth

To investigate whether the HMX3-dependent gene signatures are KMT2A::MLLT3-dependent, we performed RNA-seq analysis on CD34^+^ stem and progenitor cells (HSPC) with and without ectopically expressed *HMX3* from three unrelated donors. To this end, CD34^+^ cells were transduced with a retroviral vector expressing HMX3-T2A-GFP. As control, cells expressing GFP (GFP^+^) were used and expression of *HMX3* was confirmed by qPCR (Supplementary Figure S1I). Following purification of GFP^+^ transduced cells, RNA isolation and RNA-seq, PCA showed significant conditional gene expression changes independent of donor usage (Figure 4A). Differential expression analysis revealed 1,892 upregulated genes and 834 downregulated genes (padj < 0.05, respectively, fold change > 1.5 and < −1.5) following *HMX3* expression (Supplementary Table S4). Subsequent GSEA uncovered similar *HMX3*-expression enriched biological processes compared to primary KMT2A::MLLT3 AML cases, *i.e.* cell cycle progression and active proliferation (E2F and MYC targets, and G2M checkpoint) (Figure 4B). We conclude that *HMX3* has a major role in global gene expression regulation by controlling genes involved in cell-cycle progression and proliferation, irrespective of KMT2A::MLLT3 expression.

**Figure 4.**
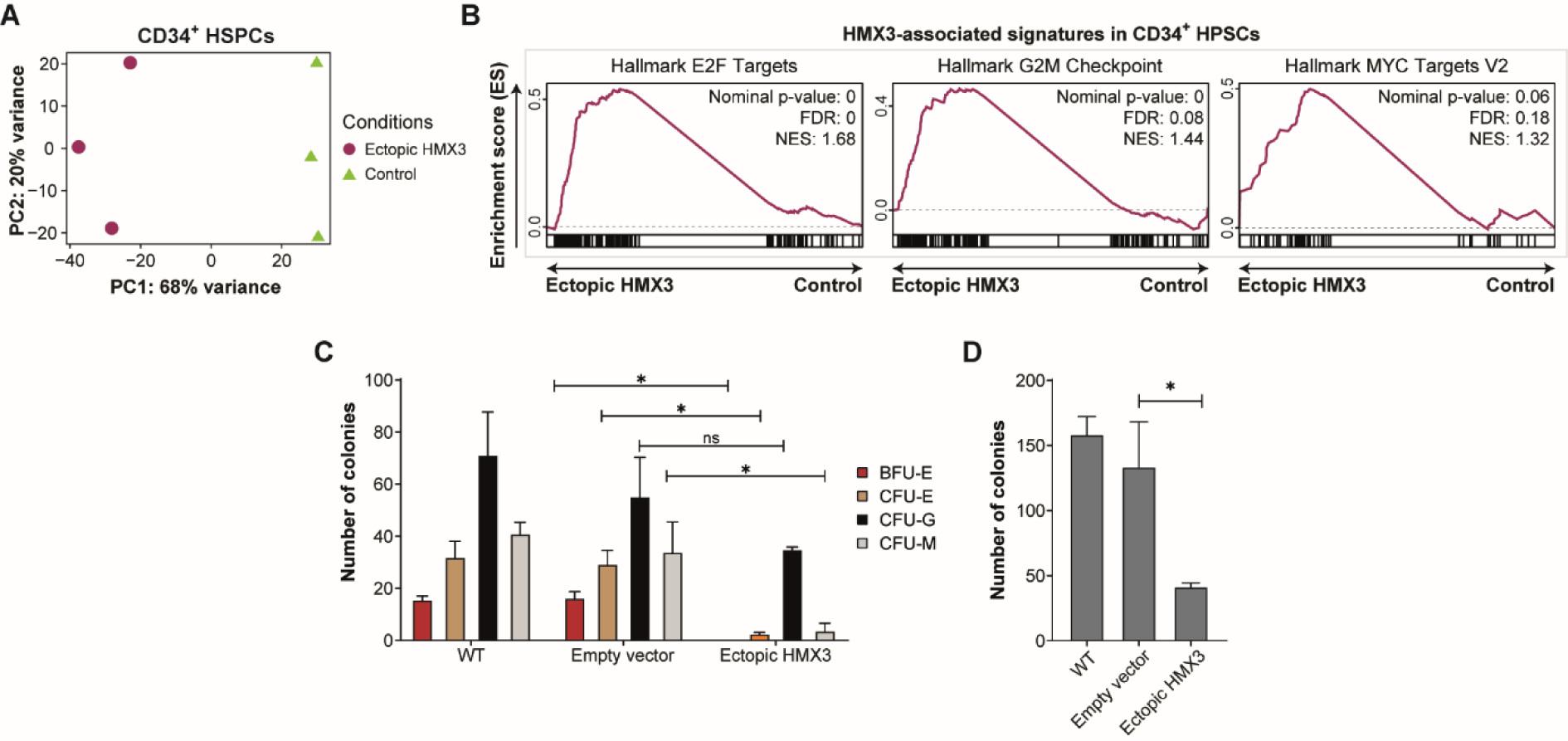
HMX3 is associated with oncogenic program in healthy CD34^+^ cells and inhibits clonogenic growth. (**A**) PCA plot using RNA-seq data from healthy CD34^+^ HSPCs transduced with an *HMX3* overexpression vector (Ectopic HMX3) and cells transduced with GFP expression (Control). Colors and shapes represent the conditions. (**B**) Enriched gene sets in healthy CD34^+^ HSPCs transduced with an *HMX3* overexpression vector (Ectopic HMX3). Colors represent the conditions from (A). (**C**) Bar chart depicting the number of colonies from CFU assays. Burst-forming unit-erythroid (BFU-E), colony-forming unit-erythroid (CFU-E), CFU-granulocyte (CFU-G), and CFU-monocyte (CFU-M) from donor-derived healthy HSPCs (WT) and HSPCs transduced with either an empty vector or with an *HMX3* overexpression vector (Ectopic HMX3). Asterisks indicate p-values < 0.05, while ‘ns’ denotes non-significant. Error bars represent standard deviation. (**D**) Bar chart representing the number of total colonies in the CFU assays. X-axis represents similar conditions as in (C). Asterisks indicate p-values < 0.05. Error bars represent standard deviation.

To determine whether HMX3 influences blood cell differentiation independent of KMT2A::MLLT3, we studied the clonogenic capacity of healthy CD34^+^ HSPCs upon forced *HMX3* expression compared to an empty vector and non-transduced cells (WT). Colony forming assays of common myeloid progenitor cells (CFU-GEMM) were performed (Figure 4C). Wild-type and empty vector-transduced CD34^+^ cells formed comparable ratios of BFU-E, CFU-E, CFU-G, and CFU-M colonies in methylcellulose culture. In contrast, we observed a significant reduction in clonogenic capacity (60-70% reduction) upon forced *HMX3* expression in the 3 independent donors (Figure 4D). While erythrocytic and monocytic colony formation was dramatically impaired upon *HMX3* expression (p < 0.01), CFU-G forming capacity was only slightly reduced (Figure 4C). Thus, HMX3 activity is permissive for granulocyte commitment but represses the formation of monocytic and erythroid colonies.

### HMX3 is a survival factor in MECOM-negative KMT2A::MLLT3 AML

To determine the oncogenic potential of HMX3, we silenced it in the two KMT2A::MLLT3 positive cell lines THP-1 and MOLM-13. Both cell lines express *HMX3* to a similar extent as primary AML samples (Supplementary Figure S1J). Following confirmation of *HMX3* silencing by qPCR (Supplementary Figure S1H and Supplementary Figure S1K-L), we performed competitive growth assays by mixing *HMX3* silenced GFP^+^ cells with non-transduced GFP^−^ cells in a 1:1 ratio (Figure 5A-B and Supplementary Figure S1M). While non-targeting transduced cells exhibited comparable expansion rates to wild type cells, *HMX3*-silenced cells (HMX3 KD) were overgrown within days. In concordance with the RNA-seq results, these data indicate that HMX3 is essential for the proliferation of KMT2A::MLLT3 cells. To determine why HMX3 KD cells were rapidly overgrown by wild type cells, we assessed the effect of HMX3 on the cell cycle by flow cytometry in MOLM-13 cells. HMX3 KD resulted in a significant increase in the percentage of cells in G0/G1 phase and decrease in the percentage of cells in S and G2/M phase (Figure 5C and Supplementary Figure S2A-B). As a cell cycle arrest may induce programmed cell death, we next determined whether HMX3 also inhibits apoptosis. To study this, we performed AnnexinV-7AAD staining using MOLM-13 cells (Figure 5D and Supplementary Figure S2C-D). At day 7 post-transduction, HMX3 KD induced a dramatic loss of viability, with up to ∼90% apoptotic cells, showing that HMX3 is a crucial factor for the survival of KMT2A::MLLT3 AML cells. Finally, as forced *HMX3* expression in healthy HSPCs impaired monocytic colony formation and KMT2A::MLLT3 AML is characterized by an arrest at the myelomonocytic stage, we determined whether *HMX3* silencing could alleviate the myelomonocytic differentiation block of MOLM-13 cells. Compared to control transduced cells, MOLM-13 HMX3 KD induced morphological changes indicative of monocyte differentiation, exemplified by indented nuclei as shown by May Grünwald Grunwald-Giemsa (MGG) staining (Figure 5E). In line with the morphological differentiation, *HMX3*-silenced MOLM-13 cells exhibited induced expression of several monocytic marker genes, such as *CD14*, *CD36*, *CD93*, *CXCL8*, and *CXCL10* (Supplementary Table S3). Thus, *HMX3* silencing triggers terminal myelomonocytic differentiation of immortalized KMT2A::MLLT3 MOLM-13 cells.

**Figure 5.**
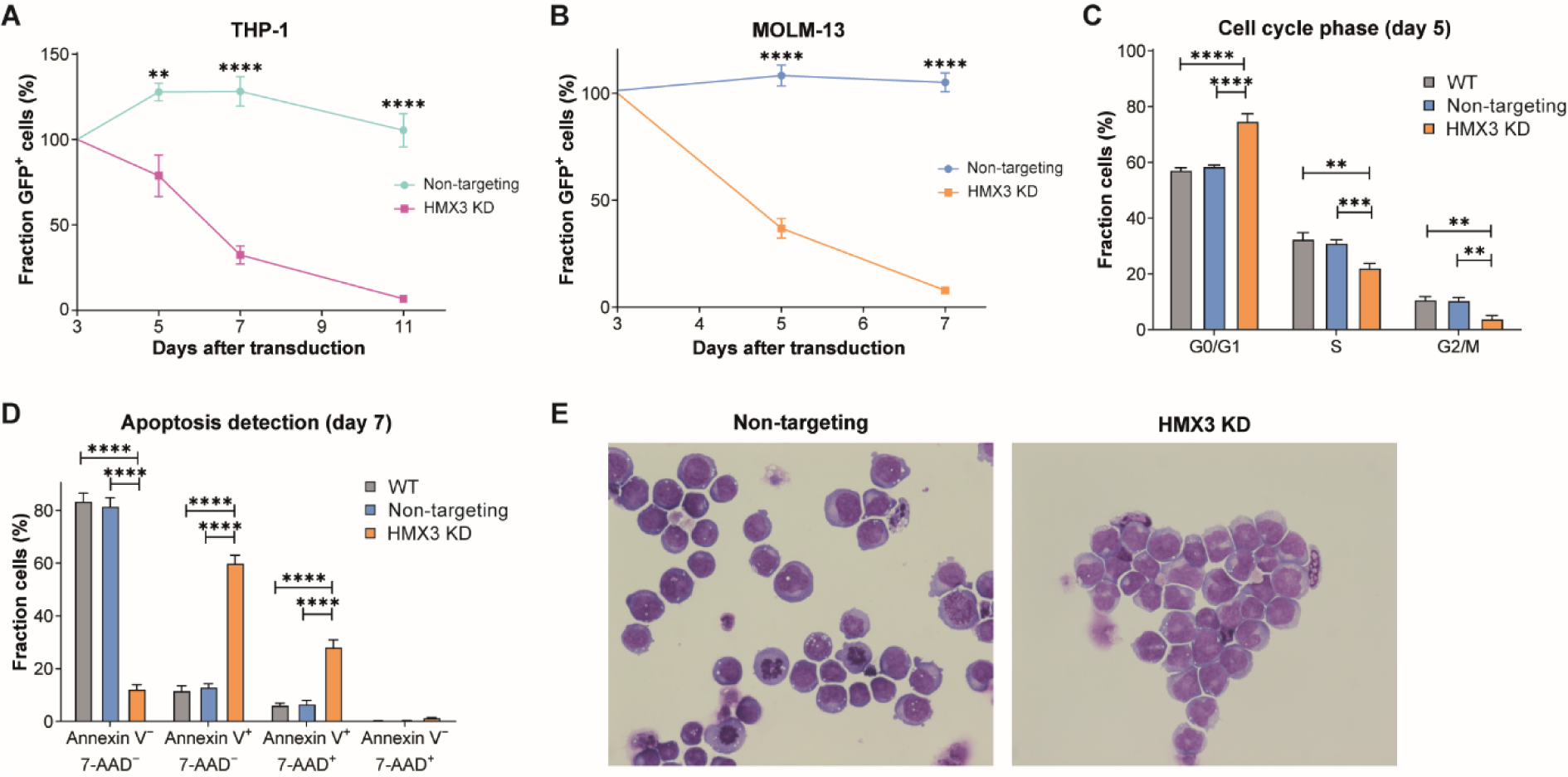
HMX3 is a survival factor for HMX3^+^ KMT2A::MLLT3 AML. (**A**) Line chart depicting competitive growth over time (days) of THP-1 cells that were either transduced with a non-targeting shRNA (Non-targeting) or a shRNA against *HMX3* (HMX3 KD), n = 3 each. Two asterisks indicate p-values < 0.01 and four indicate p-values < 0.0001. Error bars represent standard deviation. (**B**) Similar to (A), except using MOLM-13 cells. (**C**) Bar chart depicting the distribution of MOLM-13 cells across distinct cell phases (G0/G1, S, G2/M) as a percentage. Cells were measured 5 days after transduction with either non-targeting shRNA (Non-targeting) or shRNA against *HMX3* (HMX3 KD). In addition, non-transduced cells (WT) were measured. Two asterisks indicate p-values < 0.01, three indicate p-values < 0.001, and four indicate p-values < 0.0001. Error bars represent standard deviation. (**D**) Bar chart illustrating the proportion of MOLM-13 cells displaying Annexin V and/or 7-AAD positivity for assessing cell viability and apoptosis. Cells were measured 7 days after transduction with either non-targeting shRNA (Non-targeting) or shRNA against *HMX3* (HMX3 KD). In addition, non-transduced cells (WT) were measured. Three asterisks indicate p-values < 0.001 and four indicate p-values < 0.0001. Error bars represent standard deviation. (**E**) MGG staining of MOLM-13 cells 7 days after transduction with either non-targeting shRNA (Non-targeting control) or shRNA against *HMX3* (HMX3 KD) (40x magnification).

### Forced monocytic differentiation of primary pediatric KMT2A::MLLT3 AML cells following HMX3 KD

To determine whether HMX3 is crucial for the survival of primary patient cells, we silenced HMX3 in bone marrow cells taken from two independent pediatric HMX3^+^ KMT2A::MLLT3 AML cases. Following confirmation of HMX3 KD by qPCR (Supplementary Figure S2E), transduced GFP^+^ cells were used in competitive growth assays co-cultured with bone-marrow-derived mesenchymal stem cells (BM-MSC) to support expansion. Similar to MOLM-13 HMX3 KD cells, primary HMX3 KD cells were rapidly overgrown by non-transduced cells while control vector transduced AML cells were not-overgrown (Figure 6A-B). For one case, we sorted HMX3 KD and control transduced cells based on GFP-positivity followed by BM-MSC liquid co-cultures. HMX3 KD resulted in a complete inhibition of expansion while control transduced cells did expand (Figure 6C). For the other case, we sorted HMX3 KD and control transduced cells followed by RNA isolation and genome-wide RNA-seq. Compared to control cells, HMX3 KD cells showed upregulation of monocyte-associated genes, such as *CD14*, *CD300e*, *CD54*, *IL1a/b*, *IL6*, and *CXCL8* upon *HMX3* silencing (Figure 6D). In agreement with this, the proportion of CD14^+^ cells, assessed by flow cytometry, was clearly upregulated following *HMX3* KD but not in control transduced cells (Figure 6E and Supplementary Figure S2F). Finally, we determined whether HMX3 KD in primary AML cells resulted in morphological changes indicative of monocytic differentiation by MGG staining. HMX3 KD resulted in dramatically enlarged cells with indented nuclei, which is a characteristic of mature macrophages. This was not evident in control cells (Figure 6F). Together, these results indicate that HMX3 blocks the differentiation and enhances the growth of primary pediatric KMT2A::MLLT3 AML cells.

**Figure 6.**
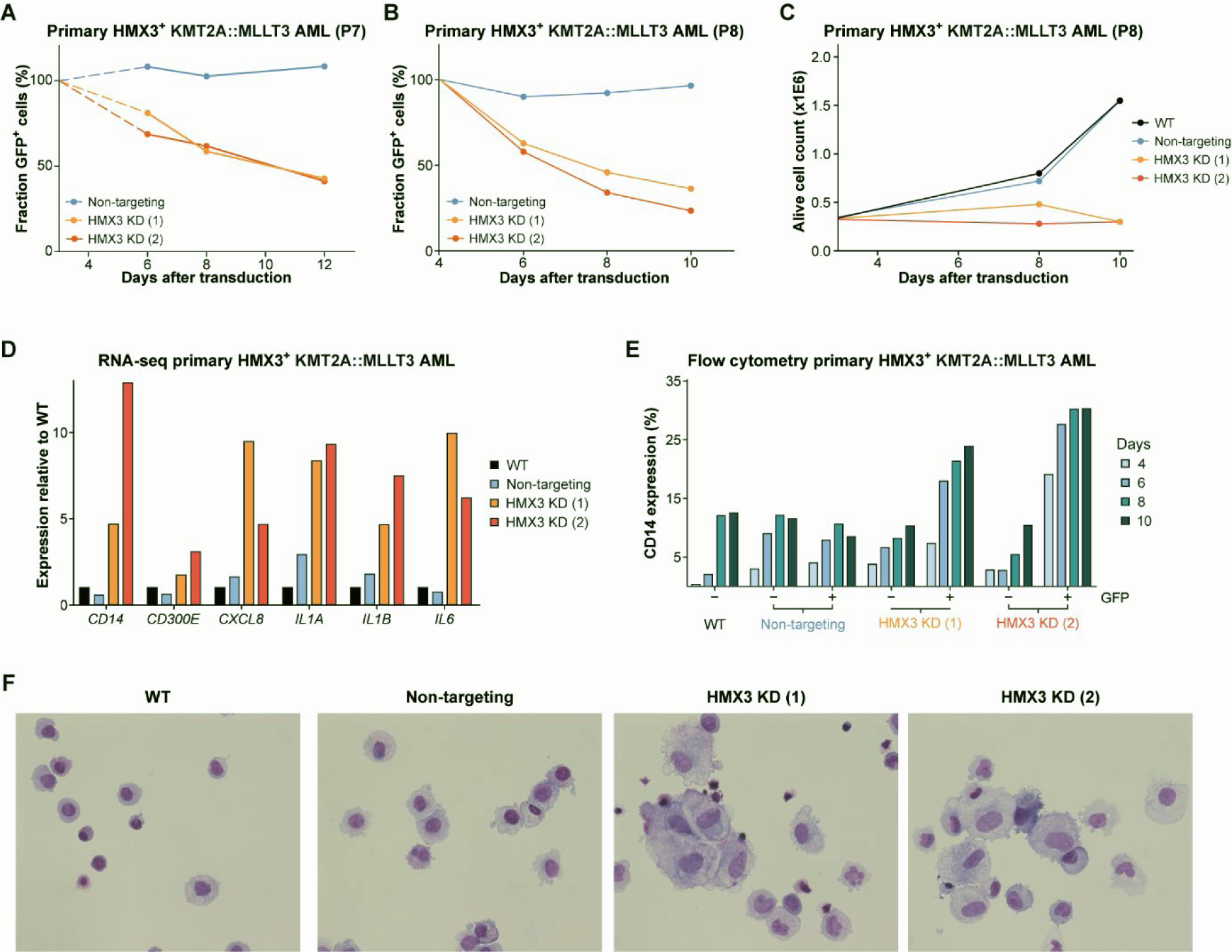
HMX3 inhibits monocytic differentiation of primary pediatric KMT2A::MLLT3 AML cells. **(A)** Line chart depicting competitive growth over time (days) of primary HMX3^+^ KMT2A::MLLT3 AML cells (P7) that were either transduced with a non-targeting shRNA control (Non-targeting) or two distinct shRNAs against *HMX3* (HMX3 KD (1) and HMX3 KD (2)). Dotted lines represent missing values. (**B**) Similar to (A), except cells from a different primary HMX3^+^ KMT2A::MLLT3 AML patient (P8) were used. (**C**) Growth curve of sorted primary HMX3^+^ KMT2A::MLLT3 AML patient (P8) after transduction with either shRNA non-targeting control (Non-targeting) or two distinct shRNAs against *HMX3* (HMX3 KD (1) and HMX3 KD (2)). In addition, wild-type non-transduced cells (WT) were measured. (**D**) Bar chart depicting normalized RNA expression relative to wild-type (WT) expression of monocytic marker genes. Conditions represent cells after transduction with either shRNA non-targeting control (Non-targeting) or two distinct shRNAs against *HMX3* (HMX3 KD (1) and HMX3 KD (2)) from AML patient P7. In addition, wild-type non-transduced cells (WT) were measured. (**E**) Bar chart representing the percentage of CD14 expressing cells measured by flow cytometry (4, 6, 8, and 10 days). Wild-type non-transduced cells (WT) of primary HMX3^+^ KMT2A::MLLT3 AML patient (P8) cells were measured, as well as cells transduced with either shRNA non-targeting control (Non-targeting) or two distinct shRNAs against *HMX3* (HMX3 KD (1) and HMX3 KD (2)). In addition, GFP positivity was measured. (**F**) MGG staining of primary HMX3^+^ KMT2A::MLLT3 AML patient (P8) cells 10 days after transduction with either non-targeting shRNA (Non-targeting control) or two distinct shRNAs against *HMX3* (HMX3 KD (1) and HMX3 KD (2)). In addition, wild-type non-transduced cells (WT) were measured.

## Discussion

### KMT2A::MLLT3 AML comes in two major subtypes

KMT2A::MLLT3 plays a crucial role in the development of acute myelomonocytic leukemia by direct activation of transforming gene programs. Here, we show that in KMT2A::MLLT3 AML the transcription factors MECOM and HMX3 are expressed in a mutually exclusive manner, where HMX3 serves as the pathological counterpart to MECOM. Just as MECOM is essential for the malignant transformation of immature myelomonocytic cells ^17,19,20^, we have established here that HMX3 plays a similarly critical role. Cases displaying MECOM expression tend to fare considerably worse in clinical outcomes and are typically of an older age ^15,16^, in contrast to HMX3-expressing cases. The latter exhibit better prognoses and are generally much younger ^15,16^. In pediatric AML in general, HMX3 is expressed in ∼70% of KMT2A::MLLT3 AML. While KMT2A::MLLT3 binds both the *MECOM* and *HMX3* locus, it is likely that this occurs in a mutually exclusive manner explaining their respective expression. Indeed, KMT2A::MLLT3 directly bound to and activated the *HMX3* gene but not the *MECOM* gene in KMT2A::MLLT3 immortalized CD34^+^ cord blood cells ^11^. In addition, we observed that the *HMX3* locus was epigenetically inactive (so very likely not bound by the activator KMT2A::MLLT3) in a MECOM^+^ KMT2A::MLLT3 AML case. These differences might be attributed by different types of stem cells being transformed in children and young adults versus older cases ^43–45^.

### HMX3 dominantly affects the transcriptome irrespective of genetic makeup

We and others observed earlier that MECOM^−^ KMT2A::MLLT3 cases share a very unique gene expression profile, while this was not evident for MECOM^+^ cases ^15,16^. As for these studies micro-array based gene expression data were used that did not include probes for *HMX3*, the expression of this TF in the MECOM^−^ group was unnoted. By performing CAGE-seq and subsequent analyses (ANANSE-CAGE) of only three AML samples of both KMT2A::MLLT3 types, we confirmed MECOM and predicted HMX3 to be key to their respective subgroups. KD studies identified several oncogenic gene programs to rely on HMX3 in KMT2A::MLLT3 MOLM-13 cells. Identical programs were identified to rely on HMX3 when primary HMX3^+^ vs MECOM^+^ KMT2A::MLLT3 were compared. Moreover, cases with the non *KMT2A*-rearranged KAT6A::CREBBP fusion also expressed *HMX3* and exhibited the unique GEP. KAT6A (a.k.a. MOZ or MYST3) activates gene expression profiles similar to KMT2A::MLLT3 ^46,47^, and both KAT6A as well as CREBBP can colocalize or interact with KMT2A ^47,48^, so it may be possible that KAT6A::CREBBP also directly activates the *HMX3* gene. Finally, very similar gene programs were found upon ectopic *HMX3* expression in normal CD34^+^ blood cells as in HMX3^+^ AML cells. Thus, HMX3 affects gene expression in a very dominant way irrespective of genetic background. It is therefore very likely that HMX3 is the driving force behind the unique GEP in genetically different types of leukemia (KAT6A::CREBBP and KMT2A::MLLT3), although few HMX3 expressing AML cases were identified that do not share the unique GEP.

### HMX3 is a leukemia-specific vulnerability in KMT2A::MLLT3 AML

Following the identification of HMX3 to be key to KMT2A::MLLT3, we showed that its knockdown in two KMT2A::MLLT3 cell lines and primary patient cells resulted in a cell cycle arrest, forced monocytic differentiation, and subsequent apoptosis. This was demonstrated based on DNA-, apoptosis- and cell surface marker staining, cell counting, gene expression analyses, and classical cell morphology. With these studies, HMX3 was identified as vulnerability similar to MECOM ^17^. MECOM has hematopoietic functions into adulthood and inherited MECOM mutations are associated with rare bone marrow failure syndrome ^49^. MECOM inhibition may therefore result in phenotypes as observed in cases with inherited MECOM mutations. While HMX3 knockout mice exhibit vestibular inner ear abnormalities and knockout females also exhibit reproductive defects, we showed here that HMX3 is not expressed in normal blood cells and HMX3 knockout mice have no discernable blood cell abnormalities ^50^. Therefore, HMX3 could serve as potential therapeutic target that once inhibited may not cause blood cell related abnormalities.

In summary, by performing CAGE-seq using a limited number of primary AML patient samples and integrating the obtained gene expression and enhancer activity data we were able to infer cell fate determining transcription factors on which the survival of both types of KMT2A::MLLT3 leukemic cells rely on. Given the functional importance of HMX3 in the pathobiology of the majority of KMT2A::MLLT3 AML and its absence in healthy blood cells, HMX3 represents a promising rationally-defined therapeutic target.

## Acknowledgements

This research was funded by Stichting Kinderen Kankervrij, project number 315. RNA-seq data results published here are part from the Beat AML study group and partly based upon data generated by The Cancer Genome Atlas managed by the NCI and NHGRI. Information about TCGA can be found at http://cancergenome.nih.gov.

## Author contributions

Conceptualization S.A.-A., B.M.H.H., B.A.v.d.R., and J.H.A.M.; methodology, S.A.-A. and S.M.B.; validation, S.A.-A. and S.M.B.; formal analysis, B.M.H.H., S.A.-A., and D.G.; investigation, S.A.-A., S.M.B., R.M., P.W.T.C.J; Resources, A.K.-H., M.V., PJ.M.V., O.H., and T.H.; writing—original draft preparation, S.A.-A.; writing—review and editing, S.A.-A., B.M.H.H., B.A.v.d.R., and J.H.A.M.; visualization, S.A.-A. and B.M.H.H.; supervision, B.A.v.d.R., J.H.J., and J.H.A.M.; funding acquisition, B.A.v.d.R. and J.H.A.M. All authors have read and agreed to the published version of the manuscript. All authors have read and agreed to the published version of the manuscript.

## STAR Methods

### Resource availability

#### Lead contact

Further information and requests for resources and reagents should be directed to and will be fulfilled by the lead contact, Bert van der Reijden.

##### Materials availability

All unique reagents generated in this study are available from the lead contact with a completed Materials Transfer Agreement.

##### Data and code availability

- RNA-seq data have been deposited at GEO and are publicly available as of the date of publication. Accession numbers are listed in the key resources table.
- This paper analyses also existing, publicly available data. The accession number for the dataset are listed in the key resources table.
- This paper does not report original code.
- Any additional information required to reanalyze the data reported in this paper is available from the lead contact upon request.

### Experimental model and study participant details

#### Primary material

All patients signed informed consent. The study was conducted in accordance with the Declaration of Helsinki and institutional guidelines and regulations from the Radboudumc Nijmegen (IRB number: CMO 2013/064). Patient data is summarized in Supplementary Table S5.

G-CSF mobilized CD34^+^ cells from healthy donors were isolated with CliniMACS CD34 Microbeads (Miltenyi Biotec, cat# 130-100-453). After 4 days of culturing in StemSpan AOF medium (Stemcell Technologies, cat# 100-0130) supplemented with 100 ng/mL rh-SCF (Immunotools, cat# 11343325), 100 ng/mL Rh-Flt3L (Immunotools, cat# 11343307), 100 ng/mL rh-IL3 (Immunotools, cat# 11340035), 100 ng/mL rh-IL6 (Immunotools, cat# 11340066), 100 ng/mL rh-TPO (Peprotech, cat# AF-300-18), and 1 µM Stem Regenin-1 (Stemcell Technologies, cat# 72342)

Mononuclear cells from KMT2A::MLLT3-positive patients (>90% blasts) were co-cultured for 3 days on a layer of primary bone-marrow-derived un-diseased mesenchymal stem cells (BM-MSC), at a density of 7500 cells/cm^2^, in StemSpan SFEM (Stemcell Technologies, cat# 9655) supplemented with 100 ng/mL rh-TPO (Peprotech, cat# AF-300-18), 150 ng/mL rh-SCF (Immunotools, cat#, 11343327), 10 ng/mL rh-Flt3L (Immunotools, cat#, 11343305), 10 ng/mL rh-IL3 (Immunotools, cat#, 11340035), 10 ng/mL rh-GM-CSF (Immunotools, cat#, 11343125), 1.35 µM UM729 (Stemcell Technologies, cat# 72332), and 750 nM Stem regenin-1 (Bio-connect, cat# c7710-1s)) before transduction (see section Transfection and transduction of cell lines) with lentiviruses containing a GFP expression construct and shRNAs against *HMX3* or non-targeting sequence on RetroNectin coated 6-well plates. After one day, cells were replated on a fresh BM-MSC feeder layer. At each timepoint (day 4, 6, 8, and 10), cells were collected for cell counts using Trypan-blue and cells were subjected to analysis on Gallios flowcytometer (Beckman Coulter) to assess expression of GFP, CD45-BV421 (Biolegend, cat# 304032), CD33-BV421 (Biolegend, cat# 303415), and CD14-APC (Biolegend, cat# 325607). On day 3, GFP-positive cells were sorted using the FACS-ARIA (BD Biosciences) for subsequent RNA isolation and further culturing on a BM-MSC feeder layer. These cells were used for cytospin preparation and May Grunwald-Giemsa staining (Merck), and cell counting at indicated timepoints.

#### Cell lines

The KMT2A::MLLT3-positive AML cell lines MOLM-13 (DSMZ, cat# ACC-554) and THP-1 (ATCC, cat# TIB-202) were used for this study. Cells were cultured at 37 °C and 5% CO_2_, and were cultured in RPMI Medium 1640 (1X) (DMEM, Gibco, cat# 21875-034) supplemented with 10% heat inactivated Fetal Calf Serum (FCS) (Biowest, cat# S1810-500). Cells were passed according to the manufacturer’s instructions. For the lentivirus production, 293FT (Gibco, cat# R700-07) cells were used. These cells were cultured at 37 °C and 5% CO_2_ in high-glucose Dulbecco’s Modified Eagle Medium (DMEM, Gibco, cat# 42430-025) supplemented with 10% non-heat inactivated FCS (Biowest, cat# S1810-500), 0.1 mM MEM Non-Essential Amino Acids (NEAA, Gibco, cat# 11140-035), 6 mM L-glutamine (Gibco, cat# 25030-024), and 1 mM MEM sodium pyruvate (Gibco, cat# 11360-070). Cells were passaged with Trypsin-EDTA (Gibco, cat# 25300-054) twice a week at a seeding density of 15%. After passing and refreshing the cells, 500 µg/µL geneticin (Gibco, cat# 11811-031) was added to the medium. For the retrovirus production, Phoenix Ampho (ATCC, CRL-3213) cells were used. These cells were cultured at 37 °C, and 5% CO_2_ in high-glucose Dulbecco’s Modified Eagle Medium (DMEM, Gibco, cat# 42430-025), supplemented with 10% heat inactivated FCS (Biowest, cat# S1810-500), and 6 mM L-glutamine (Gibco, cat# 25030-024). Cells were passaged with Trypsin-EDTA (Gibco, cat# 25300-054) twice a week at a seeding density of 15%.

## Method details

### Transfection and transduction

For the shRNA production, 293FT (Gibco, cat# R700-07) cells were grown to 30% confluence in complete medium (see cell culture section) without geneticin (Gibco, cat# 11811-031) in 9 cm dishes. 293FT cells were co-transfected using (per condition) 3 µg of pLKO.GFP transfer plasmid (Addgene, #30323) containing the shRNA (sequences in Supplementary Table S6); 4.23 µg of packaging plasmid pLP1 (Addgene, #6097) encoding GAG and POL; 1.98 µg of packaging plasmid pLP2 (Addgene, #6098) encoding REV; 2.79 µg of packaging plasmid pLP/VSVG (Addgene, #6099) encoding ENV; 62 µL 2 M CaCl2 (Sigma Aldrich, cat# C-7902), and sterile water up to 500 µL. While vortexing, 500 µL 2× HBS (HEPES-buffered saline) was added dropwise to the mixture. After an incubation of 20 minutes at room temperature, the mixture was added (1 mL) dropwise and in circles on top of the 293FT cells. After 16h of incubation, the medium was refreshed in 293FT complete culture medium without geneticin. For the virus production used in primary material experiments, the medium refreshment of the 293FT cells was done using the StemSpan SFEM II medium (Stemcell Technologies, cat# 9655). 3-4 days after transfection, virus was harvested and passed through a 0.45 µm filter. Viruses were used immediately for transduction, stored at 4 °C for 24 hours, or preserved at −80 °C after rapid freezing in liquid nitrogen. For transduction of primary cells, 20 mL of virus supernatant is concentrated using Amicon Ultra-4 30K centrifugal filters (Merck, cat# UFC803024). For transduction of suspension cells, 12 µg RetroNectin solution (TaKaRa Biomedicals, cat# T100A/B) in PBS was added to each well of untreated 6-well plates (Greiner, cat# 657185) and incubated for 2 hours at room temperature. Next, the RetroNectin solution was replaced by 2 mL of sterile 2% BSA solution (Sigma Aldrich, cat#4503) in PBS, for 30 minutes at room temperature. The BSA solution (Sigma Aldrich, cat#4503) was replaced by 2.4% v/v HEPES (Sigma Aldrich, cat#4034) on sterile PBS. The plates were stored at 4 °C. Viruses were added to the RetroNectin solution coated plates and centrifuged at 4500 xg for 1.5 hours at 4 °C. After removing the supernatant 2×105 cells/mL were added. Cells were collected 3 days post-transduction and centrifuged at 500 xg for 5 minutes at room temperature. shRNA sequencing used in this study can be found in Supplementary Table S6.

For the retrovirus production, Phoenix A cells (ATCC, cat# CRL-3213) were plated in 9 cm dishes. For each condition, 20 µL of plasmid pMIGR1 (Addgene, cat# 27490) containing the cDNA of HMX3 was combined with 62 µL of 2 M CaCl2, and then water was added to reach a total volume of 500 µL. While vortexing, 500 µL 2× HBS (HEPES-buffered saline) was added dropwise to the mixture. After an incubation of 20 minutes at room temperature, the mixture was added (1 mL) dropwise and in circles on top of the Phoenix A cells. After 16h of incubation, the medium was refreshed in Phoenix A complete culture medium. 3-4 days after transfection, virus was harvested and passed through a 0.45 µm filter. Viruses were used immediately for transduction, stored at 4 °C for 24 hours, or preserved at −80 °C after rapid freezing in liquid nitrogen. CD34^+^ cells were transduced with retroviruses on RetroNectin coated 6-well plates, similar to the cell lines

### Competitive cell growth (CCG) assay

Three days after transduction, transduced and non-transduced cells were mixed in 1:1 ratio. For this, 150 µL cell suspension was fixed using 50 µL of 4% formaldehyde. GFP was measured on the Gallios 10-color (Beckman Coulter). Dead and doublet cells were excluded based on forward and side scatter in Kaluza Analysis V2.1 software. The cells for the competitive-growth assay were plated at 0.3 million cells/mL and the GFP percentage was followed over time.

### Annexin V/7-AAD staining

The CaCl2 working solution was prepared by mixing 484.5 µL medium (see cell culture section) and 15.5 µL 2 M CaCl2 (Sigma Aldrich, cat# C-7902). This was followed by the preparation of the Annexin V solution consisting of 3 mL cell culture medium, 100 µL CaCl2 previously prepared working solution, and 2.5 µL Annexin V-Alexa Fluor 647 (Biolegend, cat# 640912), which was stored at 4 °C in the dark. Next, 3-30×104 cells were collected by centrifugation at 500 xg for 5 minutes at room temperature. Next, the cell pellet was resuspended in 150 µL of the Annexin V solution and incubated for 15 minutes at room temperature in the dark. Then, 0.5 µL 7-AAD (Sigma, cat# A9400) was added to each sample and incubated for 1 minute. Next, 50 µL of 4% formaldehyde was added and the samples were immediately measured by flow cytometer on the Beckman Coulter Gallios 10-color. Flow analysis were performed using Kaluza Analysis V2.1 software.

### Propidium iodide (PI) staining

The Propidium Iodide (PI) working solution was prepared consisting of 4.35 mL 1 g/L sodium citrate dehydrate (Merck, cat# 6448), 0.5 mL 1 mg/mL DNAse-free RNAase A (Sigma, cat# R-5503), 100 µL 1 mg/mL PI (Sigma, cat# P-4170), and 50 µL 10% Triton X-100 (Acros Organics, cat# 215682500). This solution was stored at 4 °C in the dark. Then, 30,000-300,000 cells were collected and centrifuged at 500 xg for 5 minutes at room temperature. The cell pellet was resuspended using the remaining few µL to obtain a single cell suspension. Next, 150 µL of PI working solution was added to the pellet and mixed by pipetting. After overnight incubation at 4 °C in the dark, the cells were measured on the Beckman Coulter Gallios 10-color. Cell cycle analysis was performed using Kaluza Analysis V2.1 software.

### Mass spectrometry

Cell were washed with PBS and diluted to approximately 12E6 cells per condition. After, cells were centrifuged and the pellet was lysed with 4% SDS, 100mM Tris/HCl pH 7.6, 0.1M DTT. This solution was then incubated at 95°C for 3 min. The DNA was sheared by sonication to reduce the viscosity of the sample. Next, the lysate was clarified by centrifugation at 16,000 × g for 5 min. 30 μL of protein extract was mixed with 200 μL of UA (8 M urea (Sigma, U5128) in 0.1 M Tris/HCl pH 8.5) in the filter unit and centrifugated at 14,000 × g for 15 min. After, 200 μL of UA was added to the filter unit and centrifuge at 14,000 × g for 15 min. 100 μl IAA solution (0.05 M iodoacetamide in UA) was then mixed at 600 rpm in a thermo-mixer for 1 min and incubated without mixing for 20 min. The filter units were centrifuged at 14,000 × g for 10 min. 100 μL of UA was added to the filter unit and centrifuged at 14,000 × g for 15 min. The latter two steps were repeated twice after which 100 μL of ABC (0.05 M NH4HCO3 in H2O) was added to the filter unit and centrifuged at 14,000 × g for 10 min. Again, the last two steps were repeated twice. Next, 40 μL ABC with 0.4 μg/μL trypsin (enzyme to protein ratio 1:100) was added and mixed at 600 rpm in thermo-mixer for 1 min. The units were incubated in a wet chamber at 37°C overnight. The filter units were transferred to new collection tubes and centrifuged at 14,000 × g for 10 min. After, 2× 50 μL 0.5 M NaCl was added and the solution was centrifuged at 14,000 × g for 10 min and acidified with CF3COOH. The filtrate was then desalted and purified using Stagetip. Samples were measured on a reverse-phase Easy-nLC 1000 online connected with a Thermo Q Tribrid Fusion Orbitrap. In a total of a 4 hours peptides from the experiments were separated using a 214 min gradient of acetonitrile (7% to 30%) following by washes at 60% then 95% acetonitrile. Scans were collected in data-dependent top-speed mode for 3 seconds with dynamic exclusion set at 60s.

### Cytospins and May Grunwald-Giemsa (MGG) staining

The cytospin holder was first prepared by placing the glass slide in the holder and putting the filter card and funnel on top of it. Filters were pre-wetted with 100 µL PBS supplemented with 1% FCS and 2 mM EDTA and centrifuged for 5 minutes at 45 xg. Then, 150 µL of cell suspension was added to the funnel and centrifuged again for 5 minutes at 45 xg. Finally, the cytospin holders were carefully disassembled and the glass sides were dried at room temperature overnight, whereafter the cells on the glass slides were stained with May Grunwald-Giemsa (Merck).

### Colony-forming-unit (CFU) assay

3 days after transduction of the CD34^+^ cells, 1500 GFP-positive cells were sorted on the FACS-ARIA (BD Biosciences) and collected into 1 mL of MethoCult GM H84435 (Stemcell Technologies, cat# 84435). After mixing, cells were plated in 35 mm dishes and cultured at 37 °C and 5% CO_2_. BFU-E, CFU-E, CFU-G, and CFU-M colonies were scored after 14 days.

### RNA isolation, cDNA synthesis, and RT-qPCR

Total RNA was extracted with the Nucleospin RNA Plus kit (Macheray-Nagel, cat# 740984.250). RNA was reverse-transcribed using Moloney Murine Leukemia Virus (M-MLV) Reverse Transcriptase (RT) according to the manufacturer’s instructions (Invitrogen, Thermo Fisher Scientific, cat# 28025013). For quantitative Real Time Polymerase Chain Reaction (RT-qPCR), HMX3 (Thermofisher Scientific, cat# 4351370, Hs01392772_m1) and GAPDH (Thermofisher Scientific, cat# 4352934E) primer/probe mixes were used together with Taqman 2× Universal PCR Master Mix (Thermofisher Scientific, cat# 4318157), as recommended by the manufacturer, RT-qPCR was performed in the 7500 Real Time PCR System (Applied Biosystems). The relative fold gene expression of *HMX3* was calculated with the delta-delta Ct approach, normalizing the values against the house-keeping gene *GAPDH*. Data were analyzed using 7500 Fast System Software v1.3.1 (Applied Biosystems).

### RNA sequencing

RNA was isolated using the Nucleospin RNA plus kit (Macherey-Nagel, cat# 740955.50). Library generation was performed using KAPA RNA HyperPrep Kit with RiboErase (HMR) (Roche, cat# 08098093702), with an RNA fragmentation of approximately 300 bp fragments for 6 min at 94 °C. Library size distribution was measured using High Sensitivity DNA analysis (Agilent, cat# 5067-4626) on an Agilent 2100 Bioanalyzer and its corresponding software (version B.02.08.SI648). Libraries with average sizes between approximately 300–400 bp were used for sequencing via a NextSeq 500 system (Illumina).

## Quantification and statistical analysis

### RNA sequencing analysis

The GRCh38 reference genome was indexed using STAR aligner (v2.7.9a) with Ensembl gene annotation ^52^. Bam files from the Beat AML study group were converted to paired-end fastq files using samtools (v1.12) ^38,53^. Paired-end reads were trimmed and quality was assessed using Trim Galore (v0.6.6) (https://github.com/FelixKrueger/TrimGalore). The resulting trimmed reads were mapped against the indexed reference genome in two-pass mode with the same parameters as the GDC mRNA Analysis Pipeline version Dr15plus (https://docs.gdc.cancer.gov/Data/Bioinformatics_Pipelines/Expression_mRNA_Pipeline/#rna-seq-alignment-workflow). Gene expression levels were determined using FeatureCounts (v2.0.3) with protein coding gene annotation from Ensembl GRCh38 (version 95) ^54,55^. This was then used as input for differential expression analysis using the DESeq2 package ^56^. For GSEA, DESeq2 normalized gene counts were used with GSEA (v4.2.3) ^57^. The Hallmark gene sets with 1000 permutations were used ^58^. The permutation type ‘gene_set’ was used for comparing conditions with less than 7 vs. 7 cases.

We performed fusion detection to better determine the genetic subtype of the Beat AML samples. For this, we used FusionCatcher (v1.10) with paired-end fastq files as input ^59^, STAR-Fusion (v1.7.0) ^60^, and Arriba (v1.1.0) using junction files from the STAR aligner as input ^61^. Overlap between the three fusion detection tools was determined using FuMa (v3.0.5) ^62^. For each sample, the most reliable fusion gene was selected based on both the number of tools that identified the fusion gene as well as the mean coverage of the fusion gene among the three tools.

### Cap Analysis of Gene Expression sequencing

For CAGE sequencing 2 µg of RNA with RIN>8 were sequenced by Kabushiki Kaisha DNAFORM on an Illumina system. Ribosomal RNA reads were removed from the fastq files using sortmerna (v4.3.4) ^63^. The remaining reads were aligned against Ensembl GRCh38 primary assembly reference genome using BWA-MEM (v0.7.17-r1188) ^64^. Reads with a mapping quality lower than 20 were aligned again using HISAT2 (v2.2.1) ^65^. High quality reads were merged into a single combined BAM file using samtools (v1.15.1) ^53^. Deeptools’ bamCoverage (v3.5.1) was used to make bigwig files for reads mapping on the forward and reverse strand separately ^66^.

### ANalysis Algorithm for Networks Specified by Enhancers using CAGE (ANANSE-CAGE)

Prior to TF prediction, CAGE tags were analyzed with CAGEfightR (v1.14.0) ^67^. Weakly expressing tag clusters (TCs) were discarded (>1 TPM) and TCs expressed in at least two samples were kept. Bidirectional transcription regions, that function as a proxy for enhancers, were determined using a balance threshold of 0.95 and a window size of 201 bp. Unidirectional transcription start site regions were summed per gene annotation (TxDb objects) and used for PCA and differential expression analysis, *i.e.* DESeq2 ^56^. For more details, a R markdown script is freely available at https://github.com/vanheeringen-lab/ANANSE-CAGE. CAGE tags were visualized using the UCSC Genome Browser ^68^.

Transcription factors that are key to cell fate change were determined using ANANSE (v0.4.0) ^26^. TF binding profiles were predicted using CAGE-defined bidirectional regions, motif scores, and average ReMap ChIP-seq coverage, as described previously by Heuts et al ^27^. Subsequently, gene regulatory networks (GRNs) were determined using summed unidirectional TCs per gene. Finally, influence scores, a measure for importance, were calculated using 500k edges.

### Statistical analysis and data visualization

Statistical analysis was conducted using students t-test and chi squared test, using Graphpad 5 and 9. Figures were made using Prism Graphpad 9, R studio and R (v4.0.3).

### Key resource table

| REAGENT or RESOURCE | SOURCE | IDENTIFIER |
| --- | --- | --- |
| **Antibodies** |  |  |
| CD45-BV421 | Biologend | 304032 |
| CD33-BV421 | Biologend | 303415 |
| CD14-APC | Biologend | 325607 |
| **Bacterial and virus strains** |  |  |
| **Biological samples** |  |  |
| Healthy human CD34 HSPCs | This paper | N/A |
| Primary KMT2A::MLLT3 | This paper | Table S5 |
| **Chemicals, peptides, and recombinant proteins** |  |  |
| CliniMACS CD34 Microbeads | Miltenyi Biotec | 130-100-453 |
| rh-SCF | Immunotools | 11343325 |
| Rh-Flt3L | Immunotools | 11343307 |
| rh-IL3 | Immunotools | 11340035 |
| rh-IL6 | Immunotools | 11340066 |
| rh-TPO | Peprtech | AF-300-18 |
| Stem Regenin-1 | Stemcell Technologies | 72342 |
| rh-GM-CSF | Immunotools | 11343125 |
| UM729 | Stemcell Technologies | 72332 |
| Stem regenin-1 | Bio-connect | c7710-1s |
| L-glutamine | Gibco | 25030-024 |
| geneticin | Gibco | 11811-031 |
| CaCl<sub>2</sub> | Sigma Aldrich | C-7902 |
| RetroNectin solution | TaKaRa Biomedicals | T100A/B |
| BSA solution | Sigma Aldrich | 4503 |
| Hepes | Sigma Aldrich | 4034 |
| Annexin V-Alexa Fluor 647 | Biolegend | 640912 |
| 7-AAD | Sigma | A9400 |
| sodium citrate dehydrate | Merck | 6448 |
| DNase-free RNAase A | Sigma | R-5503 |
| PI | Sigma | P-4170 |
| Triton X-100 | Acros Organics | 215682500 |
| May Grunwald-Giemsa | Merck | N/A |
| **Critical commercial assays** |  |  |
| StemSpan AOF medium | Stemcell Technologies | 100-0130 |
| StemSpan SFEM | Stemcell Technologies | 9655 |
| RPMI Medium 1640 (1X) | Gibco | 21875-034 |
| Fetal Calf Serum (FCS) | Biowest | S1810-500 |
| high-glucose Dulbecco's Modified Eagle Medium (DMEM) | Gibco | 42430-025 |
| MEM Non-Essential Amino Acids (NEAA) | Gibco | 11140-035 |
| MEM sodium pyruvate | Gibco | 11360-070 |
| Trypsin-EDTA | Gibco | 25300-054 |
| MethoCult GM H84435 | Stemcell Technologies | 84435 |
| Nucleospin RNA Plus kit | Macherey-Nagel | 740984.250 |
| Moloney Murine Leukemia Virus (M-MLV) Reverse Transcriptase (RT) | Invitrogen, Thermo Fisher Scientific | 28025013 |
| TaqMan™ Gene Expression Assay (FAM) | ThermoFisher Scientific | 4351370 |
| Taqman 2x Universal PCR Master Mix | ThermoFisher Scientific | 4318157 |
| Nucleospin RNA plus kit | Macherey-Nagel | 740955.50 |
| KAPA RNA HyperPrep Kit with RiboErase (HMR) | Roche | 08098093702 |
| High Sensitivity DNA analysis | Agilent | 5067-4626 |
| **Deposited data** |  |  |
| RNA-sequencing | This paper | To be released |
| KMT2A::MLLT3 degradation ChIP-sequencing | Olsen et al. <sup>11</sup> | GSE173599 |
| KMT2A::MLLT3 degradation RNA-sequencing | Olsen et al. <sup>11</sup> | GSE173574 |
| **Experimental models: Cell lines** |  |  |
| MOLM-13 | DSMZ | ACC-554 |
| THP-1 | ATCC | TIB-202 |
| 293FT | Gibco | R700-07 |
| Phoenix Ampho | ATCC | CRL-3213 |
| **Experimental models: Organisms/strains** |  |  |
| **Oligonucleotides** |  |  |
| shRNA | This paper | Table S6 |
| HMX3 | ThermoFisher Scientific | Hs01392772_m1 |
| Human GAPD (GAPDH) Endogenous Control | ThermoFisher Scientific | 4352934E |
| **Recombinant DNA** |  |  |
| pLKO.GFP | Addgene | 0323 |
| pLP1 | Addgene | 6097 |
| pLP2 | Addgene | 6098 |
| pLP/VSVG | Addgene | 6099 |
| pMIGR1 | Addgene | 27490 |
| **Software and algorithms** |  |  |
| STAR aligner (v2.7.9a) | Dobin et al. <sup>52</sup> | https://github.com/alexdobin/STAR/releases |
| samtools (v1.12) | Danecek et al. <sup>53</sup> | https://github.com/samtools/samtools/releases/ |
| Trim Galore (v0.6.6) | N/A | https://github.com/FelixKrueger/TrimGalore |
| FeatureCounts (v2.0.3) | Liao et al. <sup>54</sup> | https://subread.sourceforge.net/ |
| DESeq2 | Love et al. <sup>56</sup> | https://bioconductor.org/packages/release/bioc/html/DESeq2.html |
| GSEA (v4.2.3) | Subramanian et al. <sup>57</sup> | https://www.gsea-msigdb.org/gsea/index.jsp |
| FusionCatcher (v1.10) | Nicorici et al. <sup>59</sup> | https://github.com/ndaniel/fusioncatcher/releases |
| STAR-Fusion (v1.7.0) | Haas et al. <sup>60</sup> | https://github.com/STAR-Fusion/STAR-Fusion/releases |
| Arriba (v1.1.0) | Uhrig et al. <sup>61</sup> | https://github.com/suhrig/arriba/releases |
| FuMa (v3.0.5) | Watanabe et al. <sup>62</sup> | https://fuma.ctglab.nl/ |
| sortmerna (v4.3.4) | Kopylova et al. <sup>63</sup> | https://github.com/sortmerna/sortmerna/releases |
| BWA-MEM (v0.7.17-r1188) | Li et al. <sup>64</sup> | https://github.com/lh3/bwa |
| HISAT2 (v2.2.1) | Kim et al. <sup>65</sup> | https://daehwankimlab.github.io/hisat2/ |
| samtools (v1.15.1) | Danecek et al. <sup>53</sup> | https://github.com/samtools/samtools/releases/ |
| bamCoverage (v3.5.1) | Ramírez et al. <sup>66</sup> | https://deeptools.readthedocs.io/en/develop/content/tools/bamCoverage.html |
| CAGEfightR (v1.14.0) | Thodberg et al. <sup>67</sup> | https://bioconductor.org/packages/release/bioc/html/CAGEfightR.html |
| ANANSE-CAGE (v0.4.0) | Heuts et al. <sup>27</sup> | https://github.com/vanheeringen-lab/ANANSE/releases |
| Graphpad | Prism | https://www.graphpad.com/ |
| R (v4.0.3) | Posit | https://cran.r-project.org/bin/windows/base/old/ |
| R studio | R Project | https://posit.co/download/rstudio-desktop/ |
| Other |  |  |
| BeatAML Cohort data | Tyner et al. <sup>38</sup> | https://gdc.cancer.gov/about-data/publications/BEATAML1-0-COHORT-2018 |
| TARGET AML Cohort data | The Cancer Genome Atlas Program | http://cancergenome.nih.gov |

## Supplementary information

**Figure S1. Mutual exclusive *MECOM* and *HMX3* expression, ChIP-seq profiles, summary most abundant KMT2A-rearrangements, and quality control HMX3 KD in MOLM-13 and THP-1.** (**A**) Scatterplot indicating mutually exclusive *MECOM* and *HMX3* expression using our CAGE-seq samples. Expression values are normalized (TPM). Shapes indicate conditions. (**B**) UCSC genome browser illustration of HCB1 KMT2A::MLLT3-HA-FKPB12 and KMT2A::MLLT3 degraded (3h) ChIP-seq region of the HMX3 locus. In addition, RNA-seq results from MOLM-13 are depicted before and after degradation. (**C**) Scatterplot indicating mutually exclusive *MECOM* and *HMX3* expression using samples from the BeatAML cohort. Expression is RPKM normalized. (**D**) Similar to (C), except using the TARGET-AML cohort. (**E**) Venn diagram indicating the genetic abnormalities associated with *HMX3* expression using samples from the Beat AML cohort (n=21). (**F**) Similar to (E), except using the TARGET-AML cohort (n=22). (**G**) Bar chart highlighting normalized *HMX3* expression from the BLUEPRINT human hematopoietic cell RNA-seq data. Data was collected from BloodSpot and our P3 and P6 patients ^37,69^. (**H**) Bar chart representing *HMX3* expression normalized against *GAPDH*, measured by qPCR, using THP-1 cells that were either transduced with a non-targeting shRNA (Non-targeting) or a shRNA against *HMX3* (HMX3 KD (1)). Four asterisks indicate p-values < 0.0001. Error bars represent standard deviation. (**I**) Bar chart representing *HMX3* expression levels of all the colonies after CFU assay, normalized against *GAPDH*. (**J**) Bar chart representing *HMX3* expression normalized against *GAPDH*, measured by qPCR, in multiple cell models and KMT2A::MLLT3 (K::M) AML patients P7 and P8. Error bars represent standard deviation. (**K**) Bar graph depicting *HMX3* expression normalized against *GAPDH*, measured by qPCR, for THP-1 cells transduced with either a non-targeting shRNA (Non-targeting) or shRNA against *HMX3* (HMX3 KD (1)). Three asterisks indicate p-values < 0.001. Error bars represent standard deviation. (**L**) Similar to (K), except using non-targeting shRNA (Non-targeting) or shRNA against *HMX3* (HMX3 KD (2), HMX3 KD (3)). (**M**) Line chart depicting competitive growth over time (days) THP-1 cells that were either with shRNA non-targeting control (Non-targeting) or two distinct shRNAs against HMX3 (HMX3 KD (2) and HMX3 KD (3)).

**Figure S2. Flow cytometry results of cell cycle distribution, apoptosis staining and CD14 expression, and HMX3 KD validation in KMT2A::MLLT3 AML (P7).** (**A**) Bar chart depicting the distribution of MOLM-13 cells across distinct cell phases (G0/G1, S, G2/M) as a percentage. Cells were measured 3 days after transduction with either non-targeting shRNA (Non-targeting) or shRNA against *HMX3* (HMX3 KD). In addition, non-transduced cells (WT) were measured. Two asterisks indicate p-values < 0.01, three indicate p-values < 0.001, and four indicate p-values < 0.0001, while ‘ns’ denotes non-significant. Error bars represent standard deviation. (**B**) Cell-cycle distribution of MOLM-13 cells transduced with a non-targeting shRNA (Non-targeting) or *HMX3* silencing shRNA (HMX3 KD), as well as non-transduced cells (WT), 3 and 6 days after transduction. PI staining were measured by flow cytometry. (**C**) Bar chart illustrating the proportion of MOLM-13 cells displaying Annexin V and/or 7-AAD positivity for assessing cell viability and apoptosis. Cells were measured 7 days after transduction with either non-targeting shRNA (Non-targeting) or shRNA against *HMX3* (HMX3 KD). In addition, non-transduced cells (WT) were measured. One asterisk indicate p-values < 0.05 and two indicate p-values < 0.01, while ‘ns’ denotes non-significant. Error bars represent standard deviation. (**D**) Apoptosis detection of MOLM-13 cells transduced with a non-targeting shRNA (Non-targeting) or *HMX3* silencing shRNA (HMX3 KD), as well as non-transduced cells (WT), 3 and 7 days after transduction. Cells were measured by Annexin-V/7AAD apoptosis detection kit using flow cytometry. (**E**) Bar graph depicting *HMX3* expression normalized against *GAPDH*, measured by qPCR, for primary AML cells (P7) transduced with either a non-targeting shRNA (Non-targeting) or shRNA against *HMX3* (HMX3 KD (1) and HMX3 KD (2)). (**F**) CD14 measurement of primary KMT2A::MLLT3 AML cells (P8) transduced with either shRNA non-targeting control (Non-targeting) or shRNA targeting HMX3 (HMX3 KD (1) and HMX KD (3)). Cells were measured 10 days after transduction by flow cytometry.

## Supplementary tables

The following supporting information can be downloaded at: https://surfdrive.surf.nl/files/index.php/s/seKEhPXkvrhU9Zy. **Table S1**: TF prediction results using ANANE-CAGE (MECOM-negative KMT2A::MLLT3 AML versus MECOM-positive KMT2A::MLLT3 AML); **Table S2**: Mass spectrometry data MECOM+ KMT2A::MLLT3 AML versus HMX3+ KMT2A::MLLT3 AML; **Table S3**: Differential expression analysis MOLM-13 WT versus HMX3 KD; **Table S4**: Differential expression analysis CD34 HSPC ectopically expressed HMX3 versus CD34 HSC WT; **Table S5**: Patient data; **Table S6**: shRNA sequences.

