## Supplementary figures and images for "The neuronal homeobox transcription factor HMX3 is a crucial vulnerability factor in MECOM-negative KMT2A::MLLT3 acute myelomonocytic leukemia"

### Supplementary Figure S1

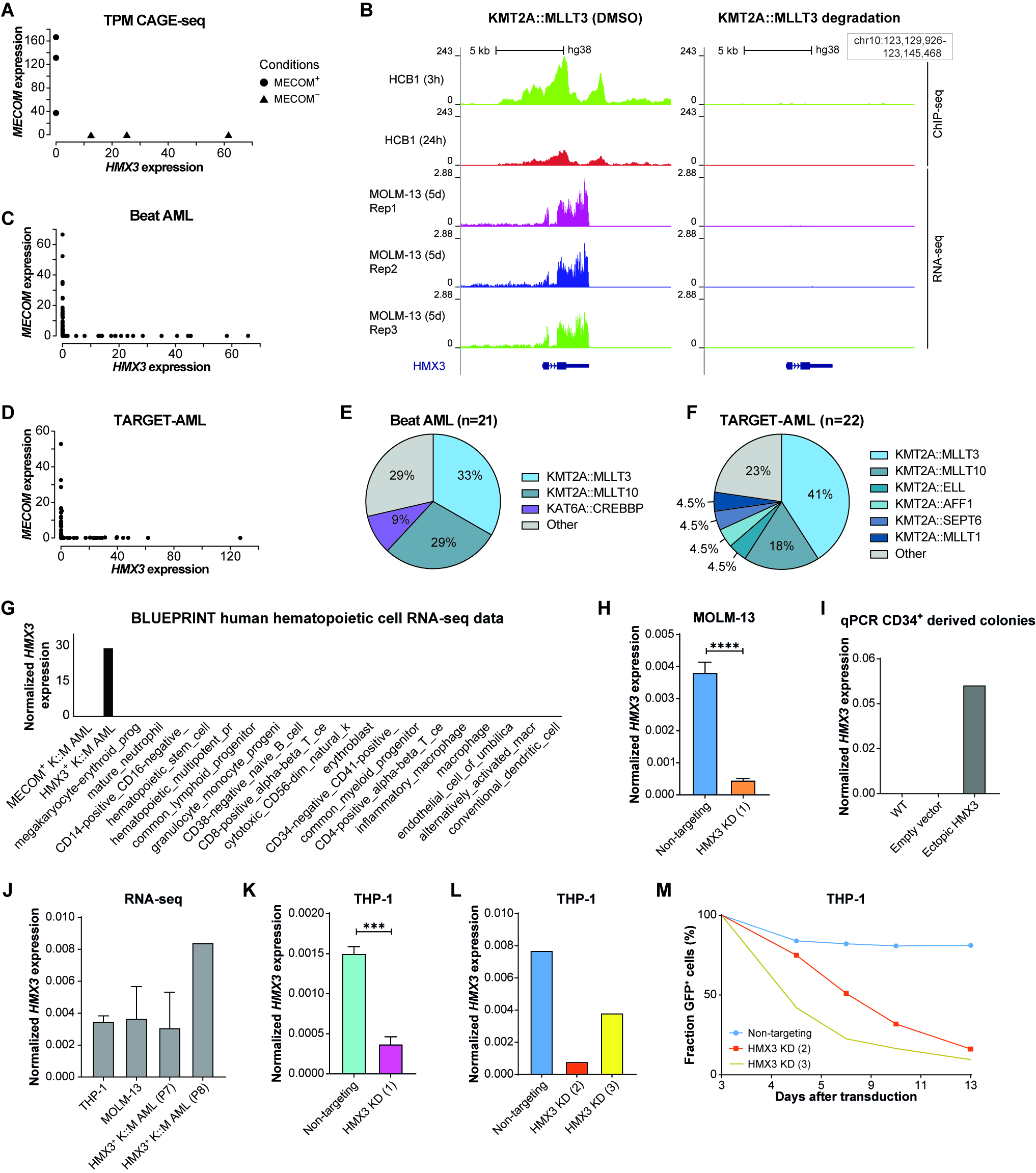

### Supplementary Figure S2

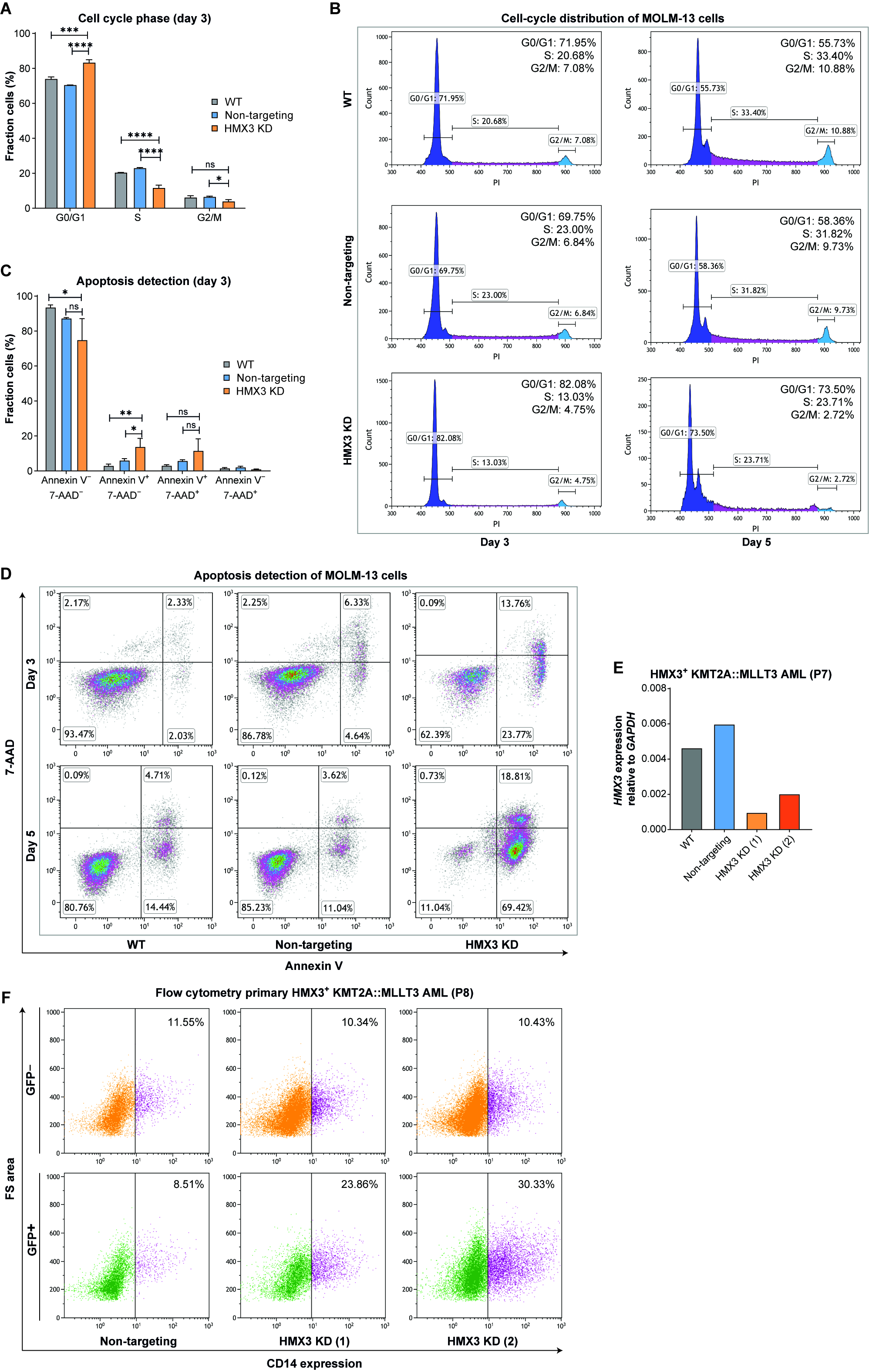
